# Major changes in plastid protein import and the origin of the Chloroplastida

**DOI:** 10.1101/799577

**Authors:** Michael Knopp, Sriram G. Garg, Maria Handrich, Sven B. Gould

## Abstract

While core components of plastid protein import (Toc and Tic) and the principle of using N-terminal targeting sequences (NTS) are conserved, lineage-specific differences are known. Rhodophytes and glaucophytes carry a conserved NTS motif, which was lost in the green lineage that also added novel proteins to Toc and Tic. Here we compare the components of plastid protein import and generated RNA-Seq, pigment profile and trans-electron microscopy data based on high-light stress from representatives of the three archaeplastidal groups. In light of plastid protein targeting, we compare the response to high-light stress of archaeplastidal representatives based on RNA-Seq, pigment profile and trans-electron microscopy data. Like land plants, the chlorophyte *Chlamydomonas reinhardtii* displays a broad respond to high-light stress, not observed to the same degree in the glaucophyte *Cyanophora paradoxa* or the rhodophyte *Porphyridium purpureum*. We find that only the green lineage encodes a conserved duplicate of the outer plastid membrane protein channel Oep80, namely Toc75 and suggest that the ability to respond to high-light stress entailed evolutionary changes in protein import, including the departure from phenylalanine-based targeting and the introduction of a green-specific Toc75 next to other import components unique to Chloroplastida. One consequence of relaxed NTS specificity was the origin of dual-targeting of plastid derived proteins to mitochondria and vice versa, using a single ambiguous NTS. Changes in the plastid protein import enabled the green lineage to import proteins at a more efficient rate, including those required for high-light stress response, a prerequisite for the colonization of land.

**High-lights:**

- Loss of Phe-based N-terminal targeting sequences (NTS) triggered the origin of dual-targeting using a single ambiguous NTS
- The Chloroplastida evolved a green-specific Toc75 for high throughput import, next to a universal and ancient Omp85 present in all Archaeplastida
- A broad response to high-light stress appears unique to Chloroplastida
- Relaxation of functional constraints allowed a broader modification of the green Toc/Tic machinery
- Critical changes in plastid targeting enabled the origin and success of the Chloroplastida and their later conquer of land

## Introduction

Mitochondria and plastids are of endosymbiotic origin and compartments surrounded by a double-membrane^1,2^. Most possess their own genomes, but the bulk of their former coding capacity was either lost or integrated into the nuclear genome^3,4^. As a consequence, most of their proteins are post-translationally imported. Guiding of precursor proteins to the matrix or stroma typically relies on N-terminal targeting sequences (NTS)^5–7^, although some exceptions are known^8–10^. Archaeplastidal plastids have a monophyletic origin^11–13^, which is also evident by the conserved nature plastid import components, a reliable indicator for the monophyly of organelles^14–16^.

While sharing a single origin, the plastids of the three algal lineages have evolved considerable differences since their divergence more than a billion years ago^17,18^. These include, but are not limited to: (i) the thickness of a remaining peptidoglycan layer^19,20^, (ii) the localisation of starch deposits^21^, (iii) the coding capacity of their genomes^3,22^, (iv) pigment composition and the types of antenna complexes used^23^, (v) the absence or presence of a xanthophyll cycle^24^ and (vi) the composition of the protein import machinery^25 26^. It raises the question to what degree the two – critical changes in protein import and changes in plastid biology– are connected, and whether one of the two conditioned or enabled the other. Though most information about plastid protein targeting stems from the green lineage^27^, several remarkable differences between the protein import in plastids of the three algal groups (Glaucophyta, Rhodopyhta, and Chloroplastida) are known.

One important difference concerns the NTS that targets proteins to the plastid stroma. Rhodophytes and glaucophytes employ a single amino acid-based motif to target proteins to their plastids^16,27–29^. In most cases this amino acid is a phenylalanine, less frequently other bulky aromatic amino acids^27,30^. The F-based motif is found at the very N-terminus of the NTS (Fig. 1) and even retained in organisms with secondary plastids of red algal origin, such as the cryptophyte *Guillardia theta*, the diatom *Phaeodactylum tricornumutm* and the parasite *Toxoplasma gondii*^31^. It is uncertain why the F-based motif was lost in Chloroplastida, but it came with several changes such as a rise in phosphorylatable serine residues that might help in avoiding erroneous targeting to the mitochondria^32,33^.

**Fig. 1:**
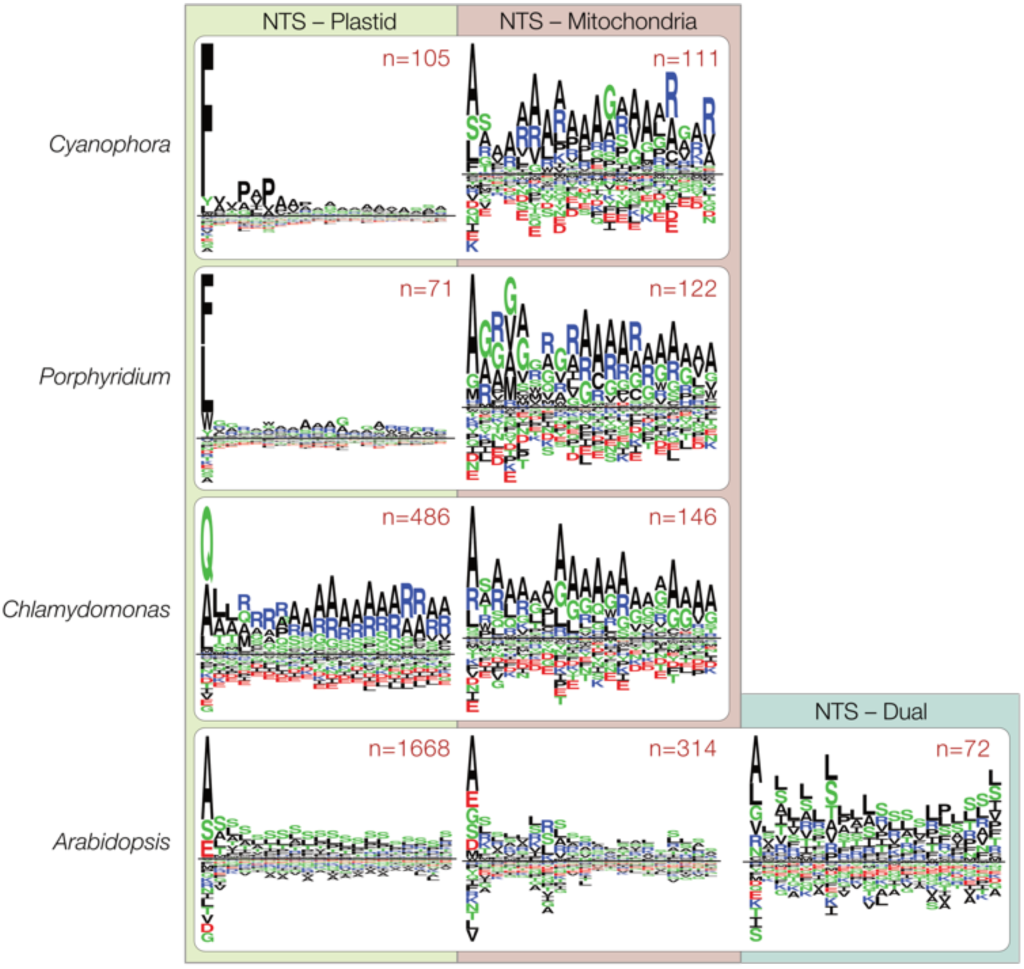
Targeting motifs and phylogenetic origin of organelle targeted proteins. **(a)** NTS of plastid- or mitochondria-targeted proteins of *C. paradoxa*, *C. merolae* and *A. thaliana*. The three species showcase the NTS for plastid- or mitochondira-targeting in the *Glaucophyta, Rhodophyta* and *Chlorophyta*, respectively. While an F-based plastid targeting motif is evident in *Glaucophyta* and *Rhodophyta*, it was lost in the green lineage.

Despite a tendency towards organelle specificity, eukaryotes also target many proteins simultaneously to two different compartments, a process known as dual-targeting. Dual-targeting can affect different combinations of compartments^34,35^, in plants also mitochondria and plastids. About 100 proteins are dually targeted to the mitochondria and plastids of *Arabidopsis thaliana* after their translation^35,36^. This large number is a consequence of the similarity between the two import mechanisms performed by Tom/Tim (translocator of the outer and inner mitochondrial membrane) and Toc/Tic (translocator of the outer and inner chloroplast membrane)^5,32^. In *A. thaliana*, a duplicate of the Toc64 receptor localizes to the outer mitochondrial membrane and now functions in mitochondrial import^37^. Both *Arabidopsis* organelles also use the same targeting-associated PURPLE ACID PHOSPHATASE2 (*At*PAP2) at their outer membranes^38,39^. The extent of dual-targeting in non-chloroplastidal species remains largely unexplored.

To investigate plastid targeting in a comparative approach across the three main algal lineages, we generated RNA-Seq, pigment profile, and trans-electron microscopy data from three different conditions (with high-light stress as the stimulus) for the chlorophyte *Chlamydomonas reinhardtii*, the rhodophyte *Porphyridium purpureum* and the glaucophyte *Cyanophora paradoxa*. The data were compared and evaluated in light of evolutionary changes regarding protein import. Our analysis connects the loss of F-based targeting and the emergence of new critical import proteins in the ancestor of the green lineage, with a series of critical changes.

## Material and Methods

### Culturing

Algae were grown in their respective media (see SAG Göttingen or ncma.bigelow.org for recipes) in aerated flasks at 20°C and illuminated with 50µE under a 12/12h day-night cycle. RNA was isolated from cells growing in the exponential phase either at 6h into the day, 6h into the night or after 1h of high-light treatment at 600µE. RNA was isolated, sequenced and assembled exactly as described previously^40^ and based on pooled biological triplicates and independently sequenced technical triplicates.

### Rapid light curves and pigment profiles

The relative electron transport rates (rETR) of the different algae were measured with use of the FluorCam FC 800MF (Photo Systems Instruments) with modulated red light (emission at 625nm and bandwidth of 40nm) as a source of measuring light (<0.1µmol quanta m^−2^ s^−1^) and modulated blue light as saturation pulse (> 8000µmol quanta m^−2^ s^−1^) (Suppl. Fig. 1). The algae samples were dark adapted for 5 min and repeatedly submitted to increasing light intensities (13, 48, 122, 160, 200, 235, 305, 375, 542, 670 µmol quanta m^−2^ s^−1^) every 11 min. The exported numeric values were fitted according to Eilers & Peeters, 1988. For each pigment extraction the pellet of 50 ml culture was resuspended with 100% acetone, homogenized and kept at −20°C over night. On the next day extracts were centrifuged and supernatant was filtered once through a 200 nm polytetrafluoroethylene membrane and then analyzed by reversed-phage high pressure liquid chromatography (HPLC) with ultraviolet/visible spectroscopy detection (Hitachi/Merck). Pigment concentrations were determined using external pigment standards isolated from spinach thylakoids^56^.

### Microscopy

For trans-electron microscopy cells were centrifuged at 800 x g and pellet was washed twice with PBS. Afterwards pellets were carefully resuspended with 2,5 % glutaraldehyde in 0,1 M cacodylate buffer and incubated for 2-3 days at 4°C. Fixed cells were then centrifuged and pellets was washed four times with 0,1 M cacodylate buffer with a minimum of 10 min incubation time and centrifugation for 2 min at 13.000 rpm. For contrasting, samples were resuspended in 2% Osmium(VIII)-oxid + 0,8% potassium hexacyanoferrate and incubated for 1 h at room temperature. Then cells were washed again five times and after addition of 3,5 % agarose and resuspension cells were incubated on ice for a minimum of 10 min until agarose became hardened. Tube tips were cut using a guillotine and the solid agar embedded pellet was pulled out and transferred to a small glass container (40 x 19 mm, 5 ml, with plastic lids). Dehydration of cell pellets was achieved using an ascending ethanol washing series starting with 60% ethanol (1 x 10 min), followed by 70% (overnight at 4°C), 80% (2 x 10 min), 90% (2 x 10 min) and 100% (1 x 10 min), finishing with 100% ethanol + molecular strainer (1 x 10 min) and propylenoxid (1 x 15 min). Afterwards epoxide resin/propylenoxid mixtures were added to the samples with increasing epoxide resin concentrations. First, a 1h incubation with epoxide resin/propylenoxid (1:2) was followed by a 1h incubation with epoxid resin/propylenoxid (1:1) and finally an overnight incubation with epoxide resin/propylenoxid (2:1) was performed. Freshly prepared epoxide resin was added the next day and samples incubated for four hours in a vacuum to remove any remaining oxygen within the epoxide resin/cell pellet solution. Finally, pellets were cut in approx. 1 mm slices with a razor blade and placed onto the tip of a notch on a rubber mat and completely covered with epoxide resin. After that epoxide resin filled mats were incubated for 24 h at 40°C followed by 24 h incubation at 60°C for complete polymerization. Probes were then cut using a ultramicroton, placed on monitoring grids and examined using trans-electron microscopy (Zeiss EM902). For the analysis of thylakoid stacks, the distances within 10 cells were counted using Fiji^57^. For each graph 10 cells were analyzed and within each cell 10 different areas counted.

### Identification of differentially expressed genes and annotation

Subsequent to the assembly via Trinity^41^ (r2013-02-25), edgeR^42^ was used to calculate the number of differentially expressed genes. The criteria for the identification were a logarithmic fold change of at least 2 and significance of 0.001 or lower. Since this approach only detected 91 differentially expressed genes for *P. purpureum*, the significance cutoff was lowered to 0.05 as suggested by the EdgeR manual (https://github.com/trinityrnaseq/trinityrnaseq/wiki). The transcripts were ranked according to mean expression values for all three light conditions and each organism. Protein annotation was performed by a BLAST search of all CDS against 112 Refseq plant and algae genomes. All BLAST hits with at least 25% local identity and a maximum E-value of 1×10^−10^ were used for annotation. In cases where the hit did not provide enough information (hypothetical proteins, predicted proteins) the next best non-hypothetical hit was selected.

### Phylogenomic analysis

The sequence dataset for the phylogenetic analysis of the Toc75/Oep80 homologs consists of 77 amino acid sequences from Chlorophytes, Rhodophytes, Cyanobacteria, Plants and one Glaucophyte. We consulted Inoue and Potter 2004 to obtain 39 amino acid sequences of Toc75 and Oep80 homologs from either the Refseq^43^ or GenBank^44^ database via their respective gene identifiers (Suppl. table 1)^45^. Additionally, 28 genomes from Chlorophytes, Rhodophytes^46^ and one Glaucophyte were downloaded either from the Refseq, GenBank or the JGI Genome Portal^47^ (Suppl. table 1). The initial set of sequences was used as query sequences to search for Toc75 and Oep80 homologs via BLASTp (version 2.5.0)^48,49^. All non-redundant hits from each subject genome with at least 25% local identity and a maximum E-value of 0.001 were added to the sequence set. Blast hits of Oep80 and Toc75 sequences of the initial sequence set were named pOep80 and pToc75 respectively.

Multiple protein sequence alignments were constructed using MAFFT (version 7.299b) with the parameters “--maxiterate 1000” and “--localpair”^50^. The initial multiple protein sequence alignment was used to check the quality of identified homologs, resulting in the removal of sequences that differed drastically in overall amino acid composition. The multiple amino acid sequence alignment was then used to construct a phylogenetic tree via RAxML^51^ (version 8.2.8) using the substitution model ‘PROTCATWAGF’ (WAG substitution Matrix and empirical base frequencies) and 100 non-parametric bootstraps. An additional tree was constructed using the new RAxML-NG with the model LG+F+R5 and 1000 bootstraps^52^ (Suppl. Fig. 2). The trees were rooted on the split between the monophyletic cyanobacterial sequences and the rest of the taxa.

Sequences of plastid-, mitochondria- and dual-targeted proteins of *A. thaliana* were obtained from Garg and Gould 2016^33^. All proteins were blasted (diamond blastp, identity cutoff: 25%, evalue cutoff: 1×10^−10^) against a database of 94 cyanobacterial and 460 alphaproteobacterial proteomes (Suppl. table 2). All hits meeting the cutoffs were plotted against all proteomes in a 2D binary heatmap. The members of each group were sorted according to phylogenetic trees from concatenated alignments, while the order of genes was determined by hierarchical clustering (hclust, method: ‘average’). The intracellular localization of the proteins was color coded.

### Identification of nuclear encoded, mitochondria- and plastid-targeted genes

Plastid-targeted proteins were identified by blasting known and manually curated plastid-targeted proteins from *A. thaliana*^33^ against the genome of *C. reinhardtii*, *C. paradoxa* and *P. purpureum* (identity of at least 50%, query coverage of at least 50%, maximum E-value of 1×10^−5^) or extracted from published proteome data when available^53,54^. To identify mitochondria-targeted proteins, we blasted all mitochondria-targeted proteins from human, mouse and rat (according to the IMPI database, marked as “Known mitochondrial”) against the genomes of the three algae (identity of at least 50%, query coverage of at least 50%, maximum E-value of 1×10^−5^). Sequence logos of the mitochondria- and plastid-targeted proteins were curated manually by aligning the first 20 amino acids following an F, Y, W or L (according to^31^) and plotted using Seq2Logo^55^.

## Results

### Adaptive changes of common photosynthetic pigments upon high-light stress

Plants react in particular to changes in light intensity^58,59^. To analyse the differences that high-light stress has on the three algae, representing the three major groups (Table 1), we set out to perform comparative studies. The algae were adapted to growing at 50 mol photons m^−2^ s^−1^ under a 12/12 day-night cycle and at 20°C. Through rapid light curves we assessed that at 600 mol photons m^−2^ s^−1^, a saturation of the photosystems was reached in all three species (Suppl. Fig. 2). For the high-light stress treatment, the algae were hence exposed to 600 mol photons m^−2^ s^−1^ for 1h. For comparison we determined the pigment profiles from cultures that were either 6h into the night or 6h into the day phase.

**Table 1:** Major differences among the three primary algae lineages and land plants, concerning their coding capacity, composition of the photosynthetic apparatus and carbon storage properties.

| Organism | Protein-coding genes |  |  | Antenna proteins | Chlorophylls | Antenna pigments | Thylakoid organization | Starch & Storage |
| --- | --- | --- | --- | --- | --- | --- | --- | --- |
|  | Nucleus | Plastid | Mitochondrion |  |  |  |  |  |
| ***Arabidopsis thaliana***<br>(*Streptophyte plant*) | 35,176 | 88 | 122 | LHC protein complex | a,b | beta-Carotin, Lutein, Neoxanthin, Violaxanthin, Antheraxanthin, Zeaxanthin | Stacked, Grana | Starch |
| ***Chara braunii***<br>(*Streptophyte algae*) | 23,546 | 105 | 46 | LHC protein complex | a,b | beta-Carotin, Lutein, Neoxanthin, Violaxanthin, Antheraxanthin, Zeaxanthin | Stacked | Starch |
| ***Chlamydomonas reinhardtii***<br>(*Chlorophyte algae*) | 14,411 | 69<sup>a</sup> | 8<sup>a</sup> | LHC protein complex | a,b | beta-Carotin, Lutein, Neoxanthin, Violaxanthin, Antheraxanthin, Zeaxanthin | Stacked | Starch |
| ***Porphyridium purpureum***<br>(*Rhodophyte*) | 8,355 | 199<sup>b</sup> | n.d. | Phycobilisomes | a | beta-Carotin, Zeaxanthin | Unstacked, equidistant and single | Glycogen, Floridean starch |
| ***Cyanophora paradoxa***<br>(*Glaucophyte*) | 27,921 (25,831) | 136<sup>a</sup> | 44<sup>s</sup> | Phycobilisomes | a | beta-Carotin, Zeaxanthin | Unstacked, equidistant and single | Floridean starch |

The glaucophyte *C. paradoxa* shows no significant change in pigment concentration or composition, neither at night nor after light stress (Fig. 2a). For the red alga *P. purpureum* we observed only very marginal changes and the concentration of pigments for the samples collected at night was the highest. Pigment concentrations seemed to slowly decrease during the day and even further under high-light stress. This was observed for all three major pigment groups at a similar rate (Fig. 2a). Only in the green alga *C. reinhardtii* the pigment composition changed significantly especially upon high-light stress (Fig. 2a). Here in particular the xanthophyll cycle, i.e. the enzyme-driven and reversible conversion of violaxanthin into zeaxanthin is evident, a component of non-photochemical quenching thought to be absent in glauco- and rhodophytes^24^. Concentrations of chlorophylls and carotenoids actually increase under high-light stress in *C. reinhardtii*, demonstrating their de novo synthesis.

**Fig. 2:**
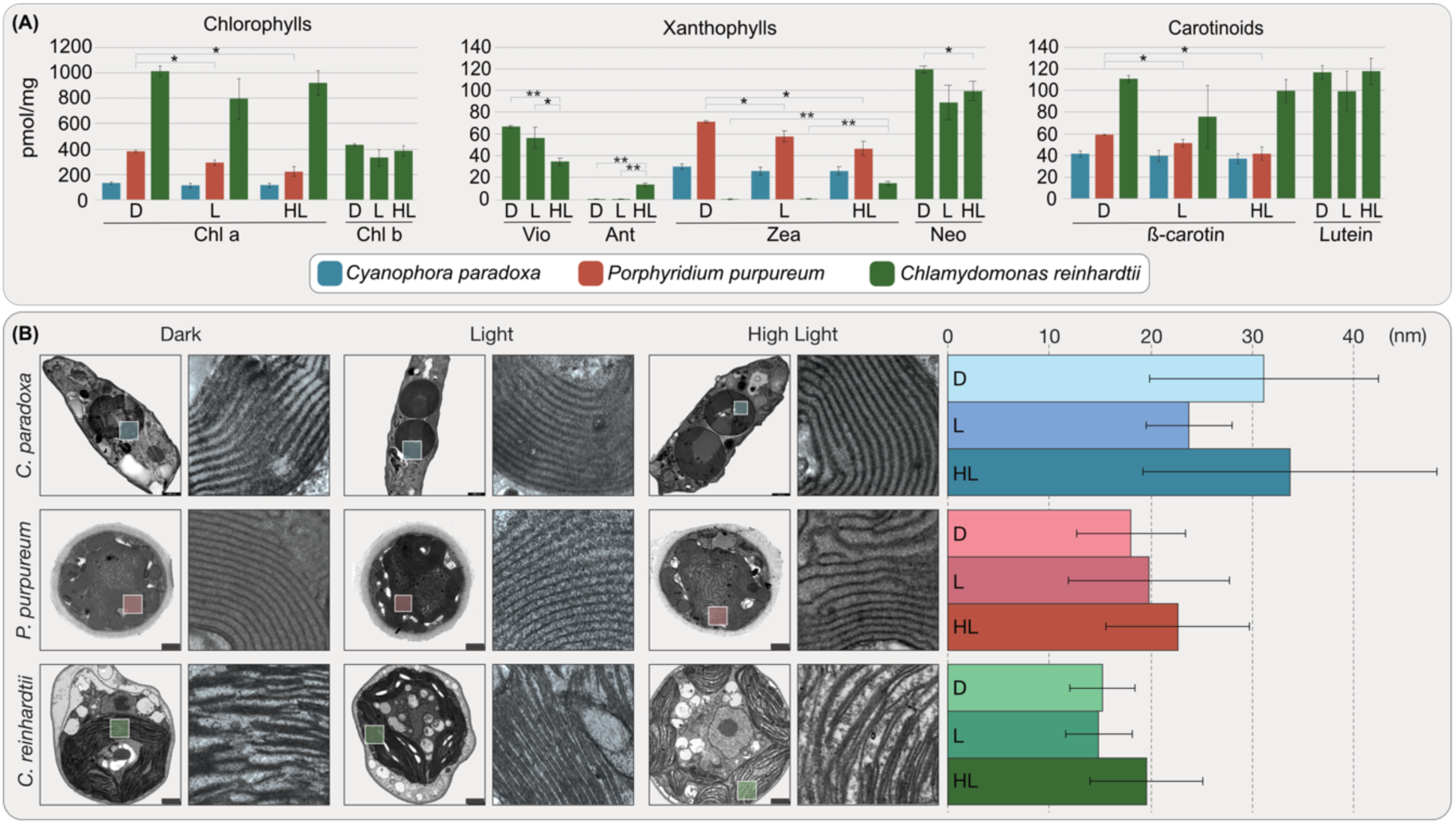
Pigment profiles and analysis of thylakoid stack distance during high-light stress. **(a)** Pigments were extracted by homogenization with acetone and their concentrations determined by an HPLC analysis. In both the glaucophyte and rhodophyte the pigment concentrations remain rather stable and only a slight decrease in the overall pigment concentration is observed during the day and even more so during high-light stress. On the contrary, in C. reinhardtii all three types of pigment change their concentration significantly and e.g. the step-wise reduction of violaxanthin (Vio) to antheraxanthin (Ant) and zeaxanthin (zea) is evident. **(b)** Cells from the three different conditions were fixed and analyzed using trans-electron microscopy and distances between the thylakoids were measured using Fiji. An obvious and statistically significant increase in thylakoid distance upon high-light stress is only observed in C. reinhardtii, although a similar but less significant trend is observed in the red alga *P. purpureum*. *<0,05 **<0,001 ***<0,0001

The thylakoid stacks (grana) of land plants relax under high-light stress in order for the repair mechanism of the photosystems to properly function^60^. This concerns in particular the degradation of the D1 protein through the membrane-bound protease FtsH, whose dimerized size is too large for the space where two thylakoid stacks align^59^. Algae form different types of thylakoid stacks^60,61^, but no grana-like structures. We performed trans-electron microscopy (TEM)-based analysis of the cells from the three different conditions and determined the distance between neighbouring thylakoid stacks. The differences we observed were in all cases marginal, but only in the case of *C. reinhardtii* did we observe a statistically significant increase in spacing upon high-light stress (Fig. 2b).

### The transcriptional response to high-light stress is most pronounced in the chlorophyte

We also generated RNA-Seq data on all samples. They reveal stark differences among the three species in terms of overall transcriptional regulation (Fig. 3). In the chlorophyte, the response to high-light stress was the most pronounced among the three algae, both regarding the number of differentially expressed genes as well as in the number of upregulated genes during high-light conditions. For each condition a clear separation was observed and a specific gene set upregulated in comparison to the average (Fig. 3).

**Fig. 3:**
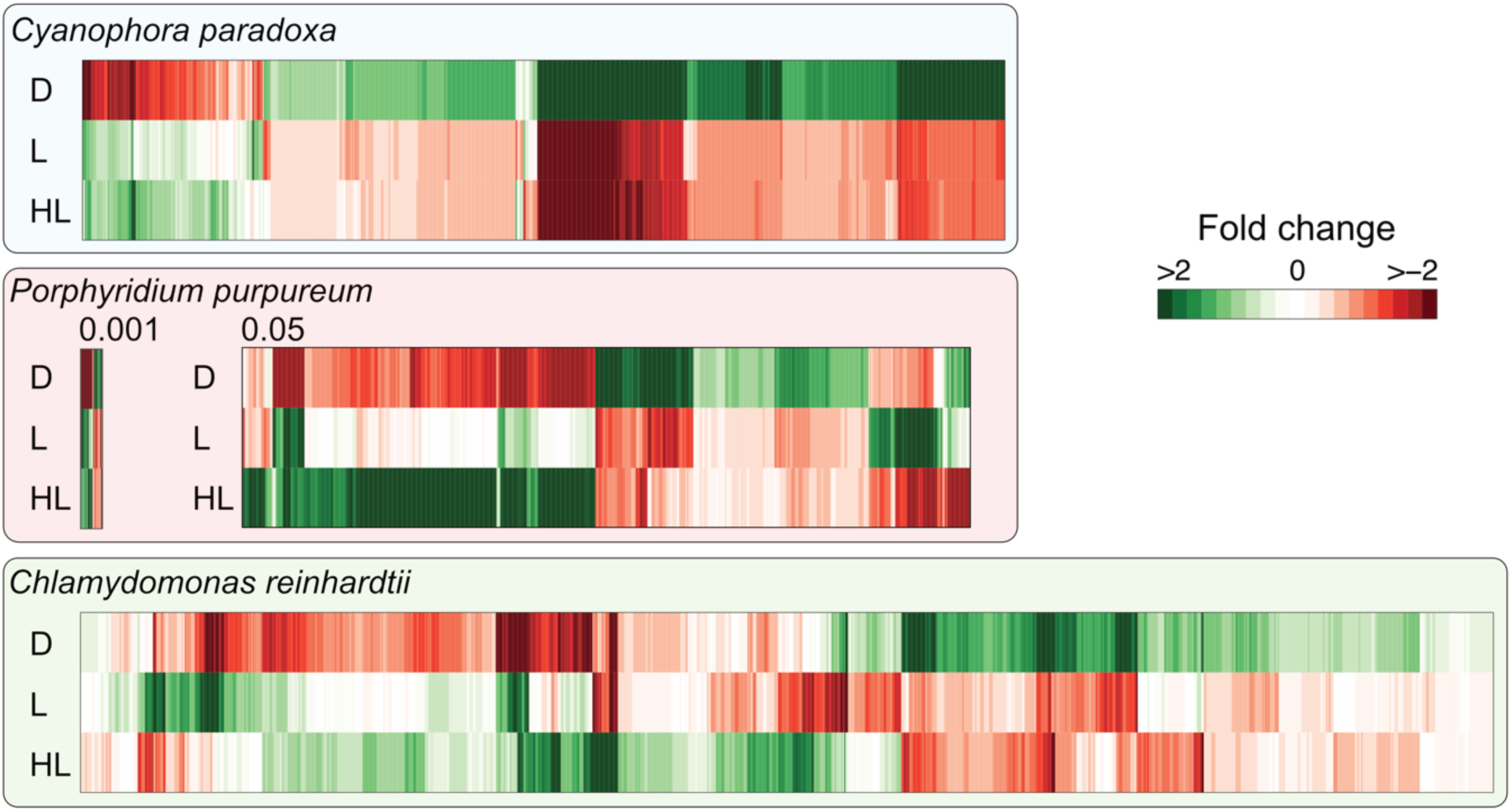
Differentially expressed genes of *C. reinhardtii*, *C. paradoxa* and *P.purpureum*. Visualization of all differentially expressed genes of *C. reinhardtii*, *C. paradoxa* and *P. purpureum*, colored according to the logarithmic fold change of the expression among all tested light conditions. Logarithmic fold changes of gene expression are color coded. For *C. reinhardtii* and *C. paradoxa*, the fold change’s significance of all visualized transcripts is at least 0.001. For *P. purpureum*, the significance cutoff was lowered to 0.05, since the original cutoff revealed only 90 differentially expressed genes. *C. reinhardtii* shows distinct sets of genes each tailored towards one of the tested light conditions. *C. paradoxa* and *P. purpureum* on the other hand, do not show such an adaptation to altering light conditions, especially not to high-light. *C. paradoxa* does not change much of its gene expression between daylight and high-light conditions, showing its lack of adaptation. Although *P. purpureum* expresses a set of genes only during high-light conditions, their differential expression was only detectable by lowering the significance cutoff. Even if all differentially expressed genes of *P. purpureum* are considered, its transcriptional changes during high-light remain minor.

Under high-light conditions the chlorophyte upregulates the expression of photosynthesis machinery components as well as proteins that promote photoprotection. A total of 418 transcripts were found to be differentially expressed, 274 values of which were significant (Suppl. table 3). The upregulated photoprotective proteins include stress-related chlorophyll binding proteins 1 and 3 involved in energy-dependent quenching to dissipate excess energy^62^, members of the early-light inducible protein family (Elip), ancestral homologs of the non-photochemical quenching associated PSBS/LHCSR3 family^65,66^, a CPD photolyase class II that reverses the formation of pyrimidine dimers that result from the exposure to strong UV radiation^67^, and chlorophyll b reductases and beta-carotene hydroxylases that prevent over-excitation of the photosystem and protects the cells from high-light intensities^68,69^. Next to these photoprotective proteins, photosynthesis house-keeping genes such as PSII Pbs27, Rieske protein, PSII subunit 28, and several proteins of the LHC superfamily were upregulated as well as a few stress-response proteins such as the plastidal homolog of DnaJ and other members of the HSP70 protein family that can form a multichaperone complex together^70^ (Suppl. table 3).

In the glaucophyte *C. paradoxa*, most of the 1,463 differentially expressed transcripts were found upregulated during darkness in correspondence to nightly proliferation. The overall difference between day and night was far more pronounced than day versus high-light and the difference between light and high-light conditions smaller than in *Chlamydomonas*. Only 26 transcripts were found to be upregulated during high-light conditions (Suppl. table 3). In comparison to the green alga, fewer proteins involved in photosynthesis regulation and photoprotection were found to be upregulated during high-light stress but also included several Elip proteins.

The identification of differentially expressed genes in *P. purpureum* posed to be more difficult. Only 90 transcripts were initially identified, but by lowering the significance cut-off — according to the standard edgeR protocol — we were able to detect another 980. The expression profile matches that of *C. paradoxa*, although the number of regulated genes is far smaller. For the transition from daylight to high-light, a total of 38 transcripts were identified (Suppl. table 3). The most notable proteins that were upregulated during high-light stress was a high-light inducible protein (Hlip) involved in non-photochemical quenching^71^, and several heat shock proteins (HSP70). The response to light stress was far weaker than in the other two algae, but *P. purpureum* might regulate its RNA levels through the extensive use of miRNAs^72^ which could contribute to the lower levels of differentially expressed genes identified.

Comparisons of the most highly upregulated proteins of each of the three algae among all conditions revealed additional differences in light-dependent differential gene expression. While *C. reinhardtii* upregulates the synthesis of several photosynthesis and plastid-related proteins during light and high-light conditions, *C. paradoxa* and *P. purpureum* upregulate only a few. In the case of *C. paradoxa*, the biggest notable difference is the focus on protein biosynthesis during darkness/night. The 50 most highly upregulated proteins during the night consist of approx. 90% ribosomal proteins, indicating an increase in overall protein biosynthesis and proliferation (Suppl. table 4). We observe photosynthesis machinery components as well as photoprotection components to be among the most upregulated proteins in combination only in *C. reinhardtii*, illustrating the chlorophyte’s more elaborate ability to adapt to differing light conditions compared to the other two screened algae.

### The red Toc75 is an Oep80, and Toc75 unique to Chloroplastida

In *Arabidopsis*, most members of its Toc75 family have been characterized. This includes the main import pore of the outer membrane, Toc75^73^ (TOC75-III, At3g46740), as well as Oep80 (TOC75-V, At5g19620) whose exact function remains unresolved while the protein is essential for plant viability^74,75^, and most recently SP2 (At3g44160) which serves protein export for chloroplast-associated protein degradation^76^. The situation in rhodophytes and glaucophytes differs and they seem not to encode the same number of Toc75 homolog^27,77^.

We collected 77 eukaryotic proteins of the Toc75 and Oep80 family from 44 eukaryotic species and routed them against their cyanobacterial homologs for the construction of a phylogenetic tree. The single glaucophyte sequence sits basal to all others, while the rhodophyte sequences form a well-supported group that is sister to all chloroplastidal sequences (Fig. 4). The sequences of green algae and plants fall into two distinct and again well-supported clusters: one comprises a group of proteins including the *At*Oep80, the other a group containing the main import pore *At*Toc75. Within these two groups separating the Oep80 from the Toc75 proteins, the separation between the chloro- and streptophytes is observed, as well as the basal branching of *Chara braunii* – a streptophyte alga related to the ancestor of land plants^78,79^ (Fig. 4).

**Fig. 4:**
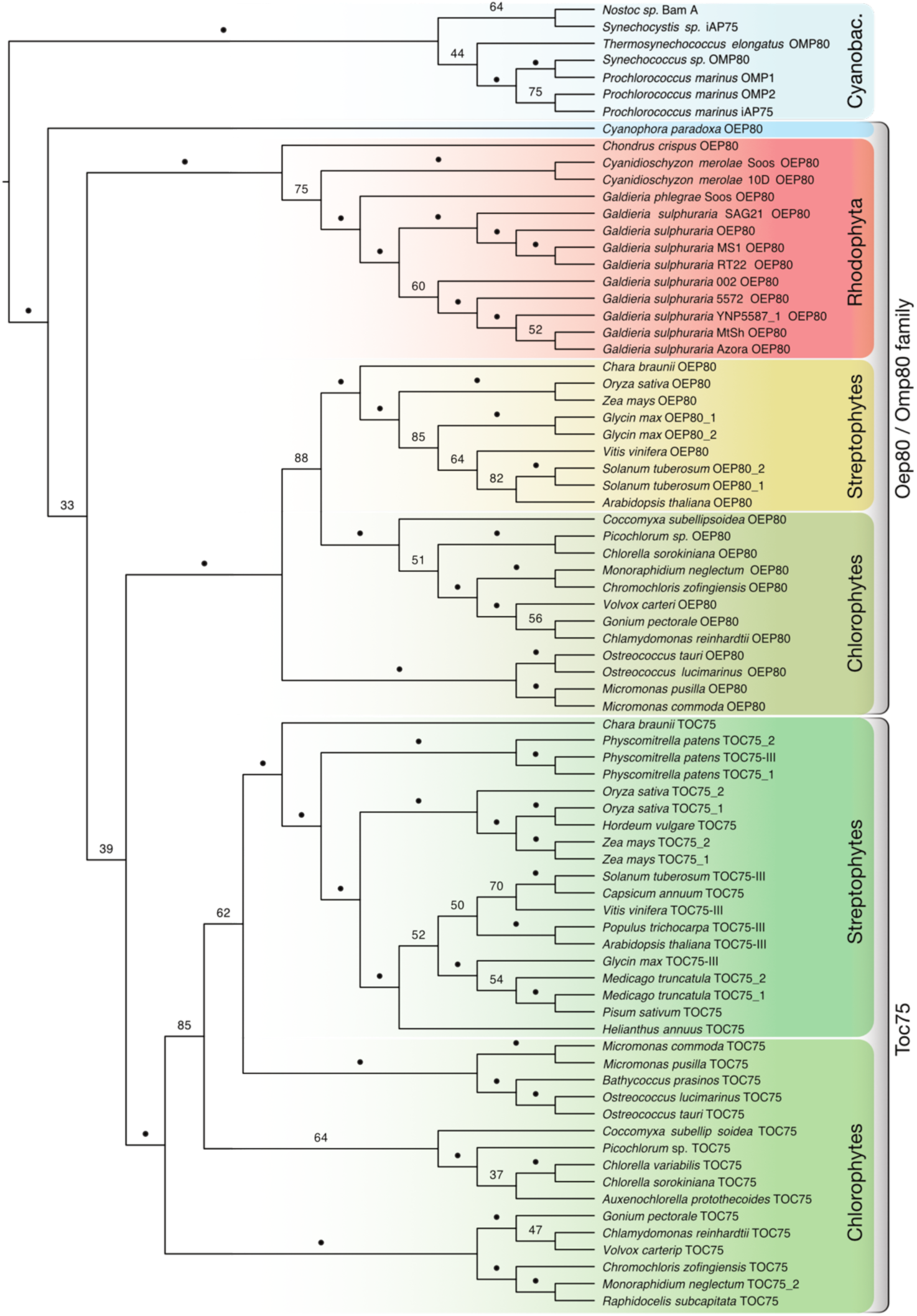
Phylogenetic analysis of Oep80 and Toc75 homologs. A total of 77 amino acid sequences of Oep80/Toc75 homologs from members of the chlorophytes, rhodophytes and glaucophytes were used for phylogeny reconstruction via RAxML (PROTCATWAGF) with 100 bootstraps. The tree was rooted on the split between the monophyletic cyanobacteria and the eukaryotic sequences. The cyanobacteria as well as all three algal groups form monophyletic groups. Within the green lineage, the Toc75 and Oep80 sequences form separate clusters, indicating the emergence of Toc75 within the green lineage.

## Discussion

If one measures evolutionary success by species diversity, the green lineage is the most successful. About 16,000 green algal, 5,000 rhodophyte and thirteen glaucophyte species have been recognized (with >100,000, 500–1000 and about a dozen that remain to be described, respectively)^80^. Another 400,000 land plant species^81^ evolved since the conquering of land some 480 million years ago^82,83^. We argue that the evolutionary origin and success of the green lineage hinges upon early changes in plastid protein targeting.

Algae and plant cells target more than a thousand proteins specifically to each of their two compartments of endosymbiotic origin. Plastid targeting evolved in a cell that had already established mitochondrial targeting, yet both import machineries share similarities and both rely on specific NTSs for matrix and stroma targeting^5^. The origin of the mitochondrial NTS is uncertain, but its positive charge was an early requirement to overcome the bioenergetic inner mitochondrial membrane^10^. The most N-terminal domain carries the charged residues critical for distinguishing between mitochondrial- and plastid targeting (Fig. 1), while the C-terminus is exchangeable^84^. Because the plastid is younger and because the photosynthetic organelle evolved in a eukaryotic cell instead of contributing to its actual origin, we understand more about the origin of the plastid NTS.

### On the origin of the N-terminal targeting sequence

It has been speculated that N-terminal targeting sequences evolved from antimicrobial peptides (AMPs)^85^, as both share similarities in terms of charged amino acid residues, the ability to form amphiphilic α-helices, and because they are frequently identified in host-endosymbiont relationships^86^. One example regarding the latter is *Paulinella chromatophora*, whose chromatophore origin is independent from that of the Archaeplastida and younger^87^. Two types of NTSs were identified that target nuclear-encoded proteins to the chromatophore, but both are not related to the simultaneously identified AMPs^88^, which argues against an AMP-origin of the NTS in *Paulinella*. The concept is also not compatible with the origin of phenylalanine-based plastid targeting and Toc75.

The components of the Toc and Tic machinery share a mixed pro- and eukaryotic ancestry^89,90^. Toc75, the β-barrel import pore in the outer membrane, is of prokaryotic origin and a member of the Omp85 superfamily^25^. Some bacterial Omp85’s recognize their substrates through a C-terminal phenylalanine^91^ and evidence is emerging that the POTRA domains of Toc75 act as binding sites for the NTS^92^. If we recall that the phenylalanine-based motif is retained in rhodophytes and glaucophytes^29^, we can conclude that the pNTS did not evolve from AMPs but rather adapted in evolution and traces back to a recognition signal for the cyanobacterial Omp85 that evolved into Toc75^93^. The ancestral character of phenylalanine-based plastid targeting was lost with the origin of the Chloroplastida and we suggest simultaneously to the expansion of the Toc75 family – with significant consequences for the green lineage.

### Dual-targeting using a single ambiguous signal is the consequence of losing the F-based motif

The use of a F-based motif offered an elegant solution to the archaeplastidal ancestor. It utilized an existing translocons-substrate recognition mechanism and allowed to distinguish cytosolically translated mitochondrial from plastid proteins through a single amino acid-based motif. With the emergence of the green Toc75 and loss of the F-based motif, false targeting likely increased. One counter-measure was the increase in phosphorylation sites in the NTS, which adds negative charge and hampers import of the substrate by mitochondria^10,34,40^. Many proteins, however, remain dually targeted in *Arabidopsis*^36^ and we predict this is restricted to the green lineage. Dual-targeting to mitochondrion and plastid does occur in algae with a red plastid, but through alternative transcription/translation initiation and not through the use of a single ambiguous NTSs^94^.

Evolution is blind. Dual-targeting evolves from falsely targeted proteins that initially might not offer a direct benefit, but are also not detrimental to the cell’s viability. This can re-localise or establish entire new pathways^95^ and there is no apparent preference regarding the direction of flow: as much proteins of cyanobacterial origin are targeted to the mitochondria as they are *vice versa* (Fig. 5). Dual-targeted proteins are largely part of the transcription and translation machinery^36^. This might include the plastid-associated polymerases, whose dual-targeting in Chloroplastida might be an ancestral trade of the lineage^79^. Both the mitochondrion and plastid have a genome, and as such information processing proteins suit a dual-targeting route well. A simultaneous control over the transcription and translation of both organelles might allow for a faster and accurate response or simply easier house-keeping. Dual-targeting reinforces the cross-communication between the two organelles of endosymbiotic origin, likely offering an evolutionary advantage to cells carrying dozens of mitochondria and plastids simultaneously such as the cells of land plants but not all algae.

**Fig. 5:**
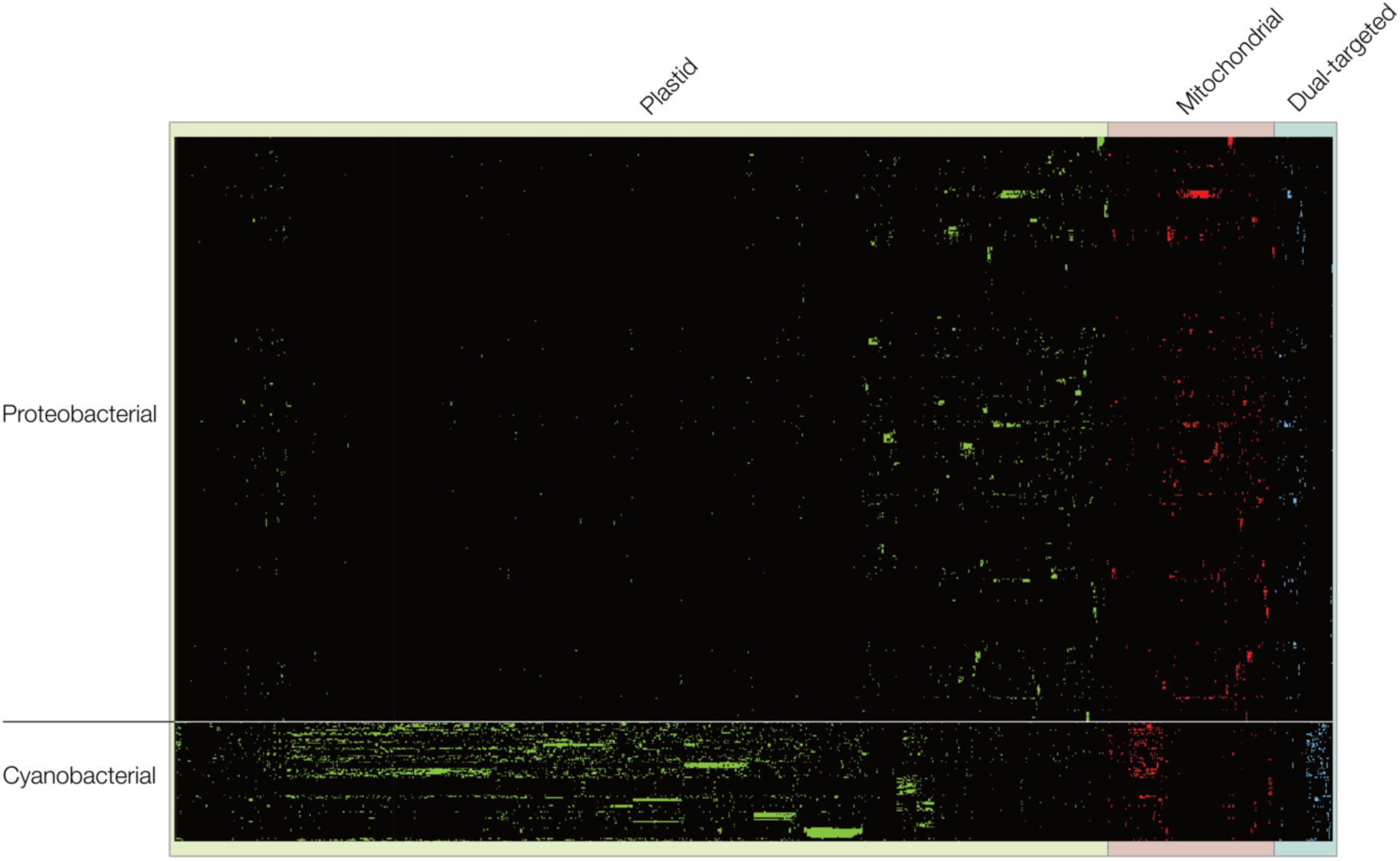
Phylogenetic oigin of plastid- and mitochondria-targeted proteins of *Arabidopsis*. Binary presence and absence pattern of homologs of plastid-(green), mitochondria-(red) and dual-targeted (blue) proteins of *A. thaliana* within 94 cyanobacterial and 460 alphaproteobacterial proteomes. Organisms are sorted according to previously constructed group-specific phylogenies, while genes are sorted by hierarchical clustering. Most homologs of plastid-targeted genes were identified in cyanobacteria, but for more than one fifth (22%) of the plastid-targeted genes the majority of homologs were identified in alphaproteobacteria. In the case of mitochondria-targeted genes, for almost one third (35%) of the genes most homologs were identified in cyanobacteria instead of alphaproteobacteria. The phylogenetic signal of the dual-targeted genes is more evenly distributed among cyanobacteria and alphaproteobacterial with one half (45%) showing a cyanobacterial origin and the other half (55%) showing an alphaproteobacterial origin.

### An Oep80 derived Toc75 is unique to the green lineage

One of the earliest descriptions of Toc75 was for a protein isolated from pea^96^. Conserved homologs across all Chloroplastida were quickly identified^15,97^, butt required way more effort across the diversity of the Archaeplastida. Through the identification of a Omp85 homolog in algae with secondary red plastids, it became evident that all phototrophic eukaryotes harbour beta-barrel forming proteins of an extended Omp85 family that form the import pore in the outer plastid membrane^98^, but with a decisive difference regarding the number of encoded homologs.

Our phylogenetic analysis of Toc75 and Oep80 supports previous analyses without the need of any sequence trimming. It demonstrates the clear-cut, likely also functional, separation between the Toc75 and Oep80 proteins of Chloroplastida^25^. The red sequences are closer to their prokaryotic homologs and the green Toc75 is further derived. From the perspective of phylogeny, there is little doubt that Toc75 is unique to the green lineage and originated from the duplication of an ancestral Omp80 that remains conserved in the other two lineages. This suggests a division of labour at the outer chloroplast membrane not found in rhodoplasts or cyanelles, the benefits of which are plenty. Glauco- and rhodophytes work with a single import pore, whereas *Arabidopsis* and its green relatives encode a single full-length Toc75 and a single full-length Oep80. Both of the latter are expressed at high levels in a conserved ratio and in the different tissues according to the gene expression atlas of the TAIR database^99^. Their presence is needed simultaneously and appears synchronized.

We speculate that the duplication of Oep80 allows for a more efficient, faster and versatile protein import. It might be a prerequisite for the elaborate response to high-light stress, which our data supports (Fig. 2, Fig. 3). A response to high-light stress is evident in all three lineages (Fig. 3), but differs in quantity and detail. *Chlamydomonas* not only alters its gene expression network the most upon high-light stress, but also focuses more on photosynthesis maintenance and protection, reacts less stressed and rapidly synthetizes pigments de-novo (Fig. 2). The upregulation of Elips that are of cyanobacterial origin occurs in all three lineages, but they were only expanded and diversified in the green lineage^100^. Retrograde signalling (a critical part of the response to high-light stress) is limited by the plastid’s import capacity^101^, highlighting the direct dependence.

If Oep80’s main duty is indeed the integration of beta-barrel proteins (and maybe other delicate substrates of unknown nature), then it releases Toc75 from this job. This mirrors the situation in mitochondria, where Tom40 acts as the main import pore while Sam50 incorporates beta-barrel proteins with a complicated topology into the outer plastid membrane^102^. The division of labour appears more effective than the simple increase in number of a single import gateway. This then maybe also allowed the endosymbiotic gene transfer of the small subunit of RubisCo to the nucleus, a trademark of the green lineage^103,104^. The sheer amount of RbcS protein required to be imported might simply overstrain the Oep80 of rhodo- and glaucophytes and its gene transfer from the plastid to the nucleus is hence selected against. These patterns allow to speculate on the sequence of evolutionary events.

Initially a duplication of the ancestral import pore Oep80 occurred and both paralogs might have performed the same duty early on. Mutations in one of the two copies led to an independence of F-based targeting, alternative substrate recognition, the emergence of NTS phosphorylation^33^, and a cytosolic 14-3-3/Hsp70-based guidance complex^105^ that we predict is unique to the green lineage, too. The plastid-encoded Tic214(YCF1)/YCF2/FtsHi complex emerged early in chlorophyte evolution, too, maybe through the duplication of an early Tic20-like protein^26,106^. The components of this complex are highly diverse, except for a C-terminal motif, and were entirely lost in grasses without impacting protein import^107,108^. Other components were added such as Tic40 that increases import efficiency^109^, and which is absent from rhodo- and glaucophytes^15^. Ever more plastid proteins went via the Toc75 route, apart from the slow folding proteins of the outer-membrane that continued to be integrated via Oep80. A more recent extension was the emergence of the CHLORAD pathway (chloroplast-associated protein degradation). Its central component, SP2, is an Omp85 paralog as well but lacks the POTRA domains^76^. It likely emerged in angiosperms and might facilitate the remodelling of plastids (e.g of a chloro-to a chromoplast), a feature unique to higher land plants and their embryoplast^18^. Therefore, the implementation of another plastid protein transport pathway based on an Oep80 duplication coincided with yet another major step in land plant evolution.

## Conclusion

Plastid endosymbiosis introduced phototrophs to the eukaryotic tree of life. A critical step was the evolution of a basic Toc/Tic protein import machinery that is conserved across all algae and plants. It is evident that major modifications of the Tic/Toc machinery and changes in the targeting sequences occurred early in the origin of the Chloroplastida. This concerns especially (i) the loss of phenylalanine-based targeting and (ii) the emergence of new import machinery components such as Tic40, a plastid-encoded Tic214, and a Toc75 that evolved from the duplication of the ancestral Omp80. We speculate that the former resulted in the emergence of dual organelle (plastid and mitochondrion) targeting using a single ambiguous targeting sequence and that the latter introduced a “high-throughput” import pathway for nuclear-encoded proteins. The main import pore of the green plastid, Toc75, is released from dealing with slow-folding proteins of the outer membrane and no longer left hamstrung when there is the need for rapid import of proteins required to cope with high-light stress. Whatever the details regarding the substrates imported by Oep80, the Chloroplastida make use of two major import pores, where rhodophytes and glaucophytes need to cope with one. Responses to high-light stress is variegated, but it requires the efficient and immediate import of over a hundred nuclear-encoded plastid proteins simultaneously after retrograde plastid signaling. This was realized by the implementation of an efficient plastid import pathway that enabled the evolutionary success of the Chloroplastida, a pinnacle of which was the conquer of land.

## Supporting information

Compiled supplementary data

## Acknowledgments

We thank Matheus Sanita Lima for discussing dual targeting and Prof. Peter Jahns for providing access to the HPLC and help in analyzing the pigment profiles. This work was supported through the DFG (267205415 – SFB 1208) and the VolkswagenStiftung (Life).

