## Supplementary material for "Major changes in plastid protein import and the origin of the Chloroplastida": Compiled supplementary data

Supplementary information:

| | |
|---|---|
| - Suppl. Fig. S1: | Rapid light curves (RLCs) of the three primary algae species |
| - Suppl. Fig. S2: | Alternative phylogenetic analysis of Oep80 and Toc75 homologs |
| - Suppl. Table 1: | Tabular listing of all sequences used to construct phylogenetic trees |
| - Suppl. Table 2: | Organisms used in the presence-absence analysis (ordered accordingly) (provided as an Excel-Table) |
| - Suppl. Table 3: | Fold changes of all differentially expressed genes between light and high light conditions (provided as an Excel-Table) |
| - Suppl. Table 4: | Fold-changes of all differentially expressed genes between all tested light conditions (provided as an Excel-Table) |
| - Suppl. Table 5: | 112 plant and algae genomes used for annotation of transcripts |

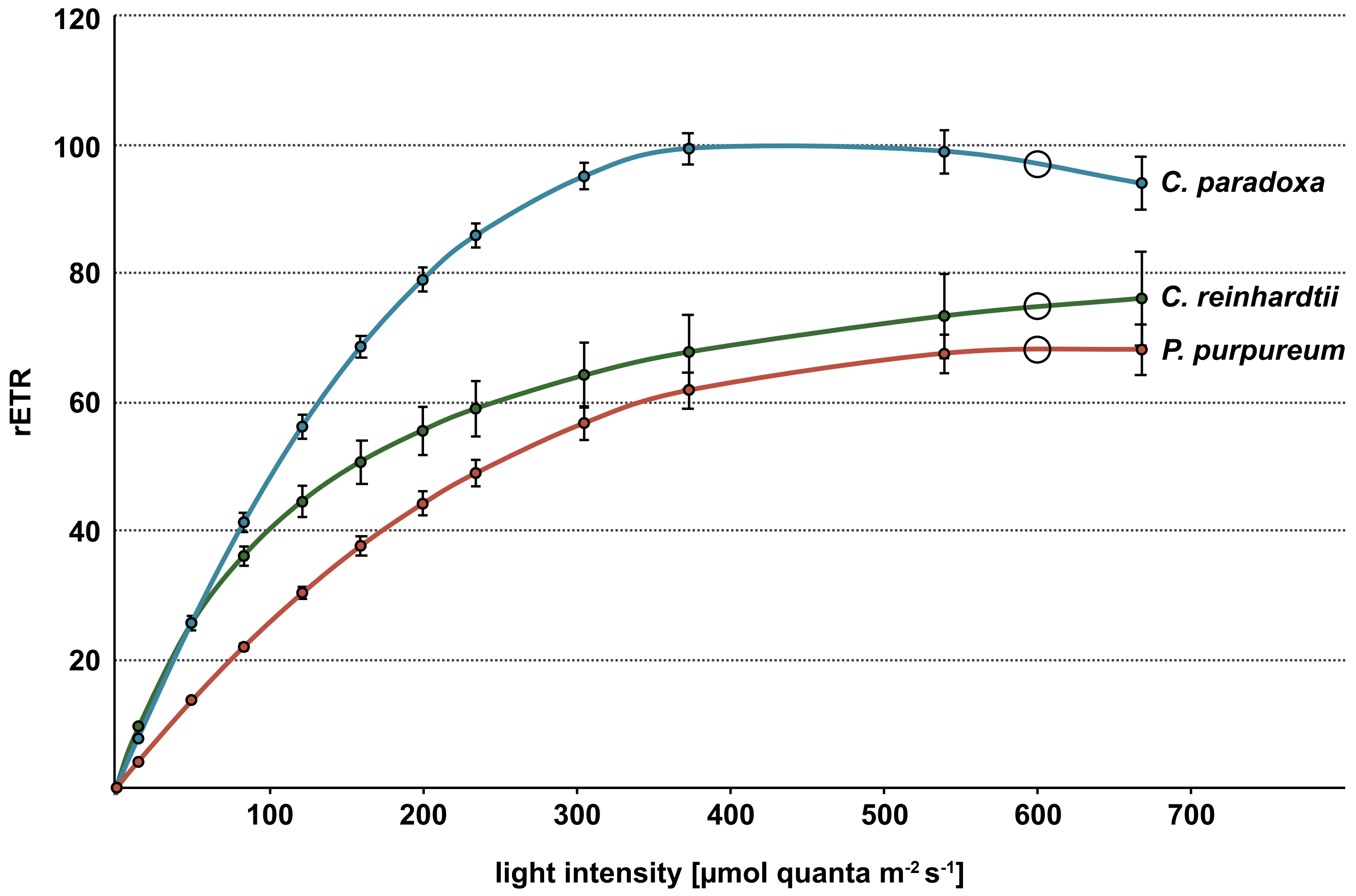


**Suppl. Fig. S1: Rapid light curves (RLCs) of the three primary algae species.** Relative electron transport rate (rETR) was determined using the FluorCam FC 800MF. Algae cultures were cultivated at 20°C with an illumination intensity of 50µE under a 12/12h day-night cycle. After 5 min of dark adaptation photosynthetic activity was measured with increasing light intensities (13, 48, 122, 160, 200, 235, 305, 375, 542, 670 µmol quanta $m^{-2}$ $s^{-1}$). At 600 µE (indicated with circles) a saturation of the photosynthetic apparatus occurred in all three lineages analyzed.

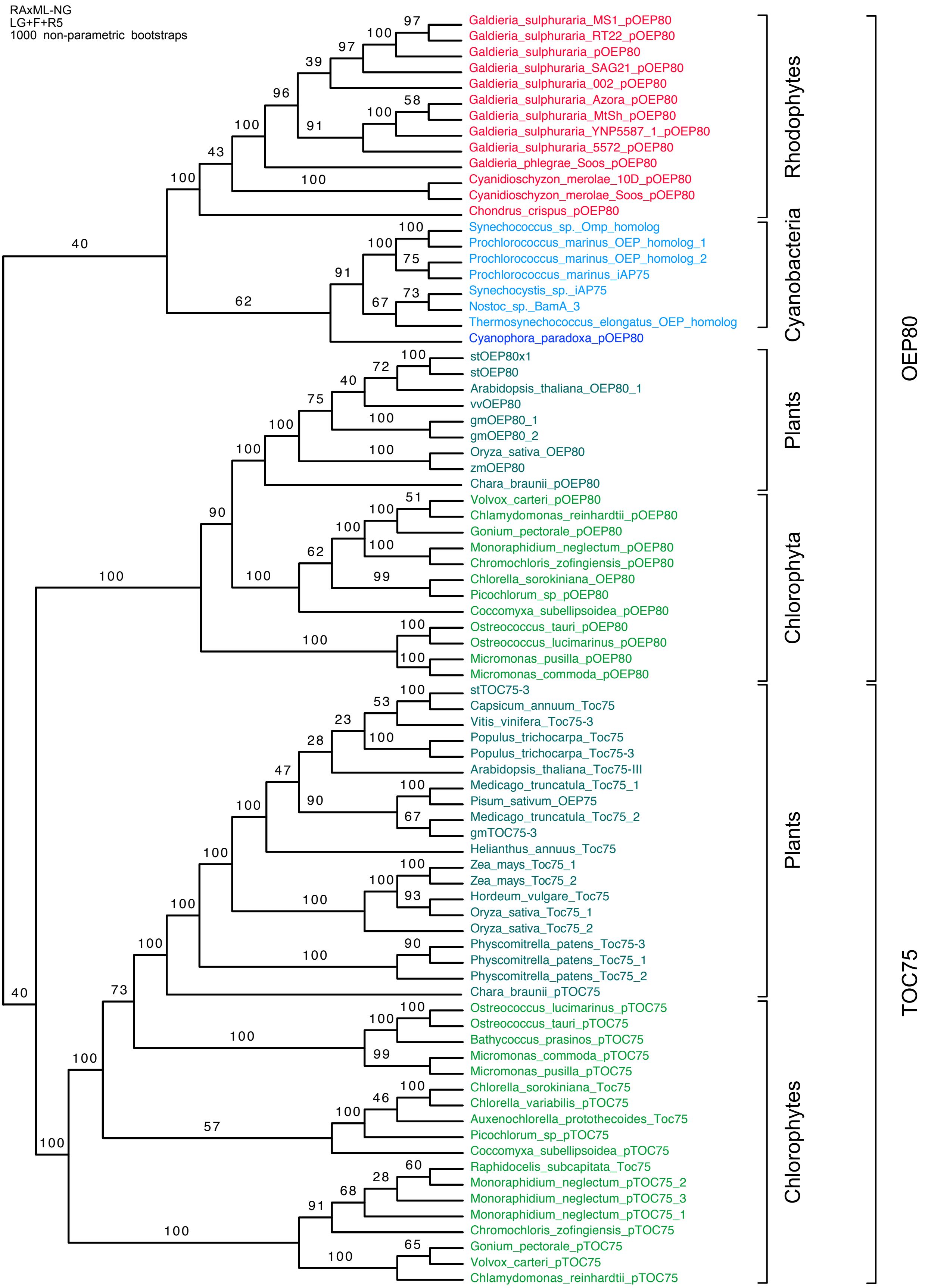


**Suppl. Fig. S2: Alternative phylogenetic analysis of Oep80 and Toc75 homologs**

Phylogeny reconstruction via RaXML-NG (LG+R5+F) with 1000 bootstraps. The tree was rooted on the split between the monophyletic cyanobacteria and the eukaryotic sequences. Although the branching pattern differs from the tree reconstructed with RAxML 8 and the PROTCATWAGF Model, all TOC75 and OEP80 sequences from *Chloroplastida* still form a monophyletic clade. Like in the analysis underlying Fig. 4, TOC75 and OEP80 proteins of plants each form monophyletic clades.

**Suppl. Table S1:** Tabular listing of all sequences used to construct phylogenetic trees (Fig. 6 and Suppl. Fig. 3) including the sequence header used in this study, accession number and taxonomic group of the source organism. All sequences that were found via BLAST searches are labelled as “Extended set”. Additionally, the genome sources of these sequences are listed including their source database, URL, download date and their respective assembly level.

| Sequence header in this study | Dataset | Group | original Accession |
|---|---|---|---|
| Bathycoccus_prasinos_pTOC75 | Extended set | Chlorophytes | XP_007509108.1 |
| Chlamydomonas_reinhardtii_pOEP80 | Extended set | Chlorophytes | XP_001695043.1 |
| Chlamydomonas_reinhardtii_pTOC75 | Extended set | Chlorophytes | XP_001703281.1 |
| Chlorella_variabilis_pTOC75 | Extended set | Chlorophytes | XP_005843553.1 |
| Chromochloris_zofingiensis_pOEP80 | Extended set | Chlorophytes | jgi\|Chrzof1\|9681\|Cz04g11310.t1 |
| Chromochloris_zofingiensis_pTOC75 | Extended set | Chlorophytes | jgi\|Chrzof1\|819\|Cz01g30030.t1 |
| Coccomyxa_subellipsoidea_pOEP80 | Extended set | Chlorophytes | XP_005648812.1 |
| Coccomyxa_subellipsoidea_pTOC75 | Extended set | Chlorophytes | XP_005649815.1 |
| Gonium_pectorale_pOEP80 | Extended set | Chlorophytes | jgi\|Gonpec1\|12926\|rna3050 |
| Gonium_pectorale_pTOC75 | Extended set | Chlorophytes | jgi\|Gonpec1\|489\|rna539 |
| Micromonas_commoda_pOEP80 | Extended set | Chlorophytes | XP_002509092.1 |
| Micromonas_commoda_pTOC75 | Extended set | Chlorophytes | XP_002504526.1 |
| Micromonas_pusilla_pOEP80 | Extended set | Chlorophytes | XP_003061676.1 |
| Micromonas_pusilla_pTOC75 | Extended set | Chlorophytes | XP_003059853.1 |
| Monoraphidium_neglectum_pOEP80 | Extended set | Chlorophytes | XP_013891338.1 |
| Monoraphidium_neglectum_pTOC75_2 | Extended set | Chlorophytes | XP_013895618.1 |
| Ostreococcus_lucimarinus_pOEP80 | Extended set | Chlorophytes | XP_001420427.1 |
| Ostreococcus_lucimarinus_pTOC75 | Extended set | Chlorophytes | XP_001416377.1 |
| Ostreococcus_tauri_pOEP80 | Extended set | Chlorophytes | XP_003081873.1 |
| Ostreococcus_tauri_pTOC75 | Extended set | Chlorophytes | XP_003074813.1 |
| Picochlorum_sp_pOEP80 | Extended set | Chlorophytes | jgi\|Picsp_1\|6034\|NSC_03388-R1_protein |
| Picochlorum_sp_pTOC75 | Extended set | Chlorophytes | jgi\|Picsp_1\|3926\|NSC_01438-R1_chloroplast |
| Volvox_carteri_pOEP80 | Extended set | Chlorophytes | XP_002948533.1 |
| Volvox_carteri_pTOC75 | Extended set | Chlorophytes | XP_002946938.1 |
| Cyanophora_paradoxa_pOEP80 | Extended set | Glaucophytes | tig00000900_g5389.t1 |
| Chondrus_crispus_pOEP80 | Extended set | Rhodophytes | XP_005711725.1 |
| Cyanidioschyzon_merolae_10D_pOEP80 | Extended set | Rhodophytes | XP_005534927.1 |
| Cyanidioschyzon_merolae_Soos_pOEP80 | Extended set | Rhodophytes | G3979.1 |
| Galdieria_phlegrae_Soos_pOEP80 | Extended set | Rhodophytes | G2108.1 |
| Galdieria_sulphuraria_002_pOEP80 | Extended set | Rhodophytes | G849.1 |
| Galdieria_sulphuraria_074W_pOEP80 | Extended set | Rhodophytes | XP_005703263.1 |
| Galdieria_sulphuraria_5572_pOEP80 | Extended set | Rhodophytes | G1359.1 |
| Galdieria_sulphuraria_Azora_pOEP80 | Extended set | Rhodophytes | G2556.1 |
| Galdieria_sulphuraria_MS1_pOEP80 | Extended set | Rhodophytes | G2257.1 |
| Galdieria_sulphuraria_MtSh_pOEP80 | Extended set | Rhodophytes | G1270.1 |
| Galdieria_sulphuraria_RT22_pOEP80 | Extended set | Rhodophytes | G2280.1 |
| Galdieria_sulphuraria_SAG21_pOEP80 | Extended set | Rhodophytes | G2095.1 |
| Galdieria_sulphuraria_YNP5587_1_pOEP80 | Extended set | Rhodophytes | G925.1 |
| Auxenochlorella_protothecoides_Toc75 | Initial sequence set | Chlorophytes | KFM25192.1 |
| Chlorella_sorokiniana_OEP80 | Initial sequence set | Chlorophytes | PRW60585.1 |
| Chlorella_sorokiniana_Toc75 | Initial sequence set | Chlorophytes | PRW61091.1 |
| Raphidocelis_subcapitata_Toc75 | Initial sequence set | Chlorophytes | GBF94877.1 |
| Nostoc_sp._BamA_3 | Initial sequence set | Cyanobacteria | BAB73968.1 |
| Prochlorococcus_marinus_iAP75 | Initial sequence set | Cyanobacteria | AAQ00463.1 |
| Prochlorococcus_marinus_OEP80_1 | Initial sequence set | Cyanobacteria | CAE19797.1 |
| Prochlorococcus_marinus_OEP80_2 | Initial sequence set | Cyanobacteria | CAE21588.1 |
| Synechococcus_sp._Omp_homolog | Initial sequence set | Cyanobacteria | CAE07070.1 |
| Synechocystis_sp._iAP75 | Initial sequence set | Cyanobacteria | BAA17512.1 |
| Thermosynechococcus_elongatus_OEP_homolog | Initial sequence set | Cyanobacteria | BAC09341.1 |
| Arabidopsis_thaliana_OEP80_1 | Initial sequence set | Streptophytes | NP_568378.1 |
| Arabidopsis_thaliana_Toc75-III | Initial sequence set | Streptophytes | Q9STE8.1 |
| Capsicum_annuum_Toc75 | Initial sequence set | Streptophytes | PHT80225.1 |
| Chara_braunii_pOEP80 | Initial sequence set | Streptophytes | GBG89700.1 |
| Chara_braunii_pTOC75 | Initial sequence set | Streptophytes | GBG61930.1 |
| Glycin_max_OEP80_1 | Initial sequence set | Streptophytes | XP_003542049.2 |
| Glycin_max_OEP80_2 | Initial sequence set | Streptophytes | XP_003547118.1 |
| Glycin_max_TOC75-3 | Initial sequence set | Streptophytes | XP_003547008.1 |
| Helianthus_annuus_Toc75 | Initial sequence set | Streptophytes | OTG28259.1 |
| Hordeum_vulgare_Toc75 | Initial sequence set | Streptophytes | BAJ97575.1 |
| Medicago_truncatula_Toc75_1 | Initial sequence set | Streptophytes | XP_003606719.1 |
| Medicago_truncatula_Toc75_2 | Initial sequence set | Streptophytes | XP_003597400.3 |
| Oryza_sativa_OEP80 | Initial sequence set | Streptophytes | XP_015627644.1 |
| Oryza_sativa_Toc75_1 | Initial sequence set | Streptophytes | XP_015630560.1 |
| Oryza_sativa_Toc75_2 | Initial sequence set | Streptophytes | XP_025877151.1 |
| Physcomitrella_patens_Toc75_1 | Initial sequence set | Streptophytes | XP_024357726.1 |
| Physcomitrella_patens_Toc75_2 | Initial sequence set | Streptophytes | XP_024357726.1 |
| Physcomitrella_patens_Toc75-3 | Initial sequence set | Streptophytes | XP_024357726.1 |
| Pisum_sativum_OEP75 | Initial sequence set | Streptophytes | CAA58720.1 |
| Populus_trichocarpa_Toc75-3 | Initial sequence set | Streptophytes | XP_002303729.2 |
| Solanum_tuberosum_OEP80_1 | Initial sequence set | Streptophytes | XP_006354253.1 |
| Solanum_tuberosum_OEP80_2 | Initial sequence set | Streptophytes | XP_006351245.1 |
| Solanum_tuberosum_TOC75-3 | Initial sequence set | Streptophytes | XP_006350787.1 |
| Vitis_vinifera_OEP80 | Initial sequence set | Streptophytes | XP_002285507.2 |
| Vitis_vinifera_Toc75-3 | Initial sequence set | Streptophytes | XP_002280661.1 |
| Zea_mays_OEP80 | Initial sequence set | Streptophytes | XP_008645435.1 |
| Zea_mays_Toc75_1 | Initial sequence set | Streptophytes | PWZ07718.1 |
| Zea_mays_Toc75_2 | Initial sequence set | Streptophytes | PWZ53262.1 |

| Organism | Source | Genome URL | Downloaded | Assembly level |
|---|---|---|---|---|
| Ostreococcus tauri | Refseq | ftp://ftp.ncbi.nlm.nih.gov/genomes/refseq/plant/Ostreococcus_tauri/latest_assembly_versions/GCF_000214015.3_version_140606 | 20.03.19 | chromosome |
| Chlamydomonas reinhardtii | Refseq | ftp://ftp.ncbi.nlm.nih.gov/genomes/refseq/plant/Chlamydomonas_reinhardtii/latest_assembly_versions/GCF_000002595.1_v3.0 | 20.03.19 | scaffold |
| Micromonas commoda | Refseq | ftp://ftp.ncbi.nlm.nih.gov/genomes/refseq/plant/Micromonas_commoda/latest_assembly_versions/GCF_000090985.2_ASM9098v2 | 20.03.19 | complete genome |
| Micromonas pusilla | Refseq | ftp://ftp.ncbi.nlm.nih.gov/genomes/refseq/plant/Micromonas_pusilla/latest_assembly_versions/GCF_000151265.2_Micromonas_pusilla_CCMP1545_v2.0 | 20.03.19 | scaffold |
| Ostreococcus lucimarinus | Refseq | ftp://ftp.ncbi.nlm.nih.gov/genomes/refseq/plant/Ostreococcus_sp._lucimarinus_/latest_assembly_versions/GCF_000092065.1_ASM9206v1 | 20.03.19 | complete genome |
| Chlorellavariabilis | Refseq | ftp://ftp.ncbi.nlm.nih.gov/genomes/refseq/plant/Chlorella_variabilis/latest_assembly_versions/GCF_000147415.1_v_1.0 | 20.03.19 | scaffold |
| Volvoxcarteri | Refseq | ftp://ftp.ncbi.nlm.nih.gov/genomes/refseq/plant/Volvox_carteri/latest_assembly_versions/GCF_000143455.1_v1.0 | 20.03.19 | scaffold |
| Bathycoccus prasinos | Refseq | ftp://ftp.ncbi.nlm.nih.gov/genomes/refseq/plant/Bathycoccus_prasinos/latest_assembly_versions/GCF_002220235.1_ASM222023v1 | 20.03.19 | chromosome |
| Coccomyxa subellipsoidea | Refseq | ftp://ftp.ncbi.nlm.nih.gov/genomes/refseq/plant/Coccomyxa_subellipsoidea/latest_assembly_versions/GCF_000258705.1_Coccomyxa_subellipsoidae_v2.0 | 20.03.19 | contig |
| Monoraphidium neglectum | Refseq | ftp://ftp.ncbi.nlm.nih.gov/genomes/refseq/plant/Monoraphidium_neglectum/latest_assembly_versions/GCF_000611645.1_mono_v1 | 20.03.19 | scaffold |
| Gonium pectorale | JGI | https://genome.jgi.doe.gov/portal/pages/dynamicOrganismDownload.jsf?organism=Gonpec1 | 20.03.19 | – |
| Chromochloris zofingiensis | JGI | https://genome.jgi.doe.gov/portal/pages/dynamicOrganismDownload.jsf?organism=Chrzof1 | 20.03.19 | – |
| Picochlorum sp. | JGI | https://genome.jgi.doe.gov/portal/pages/dynamicOrganismDownload.jsf?organism=Picsp_1 | 20.03.19 | – |
| Cyanophora paradoxa | Institute of Plant Biochemistry, HHU,Germany | http://cyanophora.rutgers.edu/cyanophora/Cyanophora_paradoxa_MAKER_gene_predictions-022111-aa.fasta | 20.03.19 | contig |
| Galdieria sulphuraria 074W | Refseq | ftp://ftp.ncbi.nlm.nih.gov/genomes/refseq/plant/Galdieria_sulphuraria/latest_assembly_versions/GCF_000341285.1_ASM34128v1 | 20.03.19 | scaffold |
| Galdieria phlegrae Soos | Institute of Plant Biochemistry, HHU,Germany | http://porphyra.rutgers.edu/Rossoni_et_al_2019.zip | – | – |
| Galdieria sulphuraria 002 | Institute of Plant Biochemistry, HHU,Germany | http://porphyra.rutgers.edu/Rossoni_et_al_2019.zip | – | – |
| Galdieria sulphuraria 5572 | Institute of Plant Biochemistry, HHU,Germany | http://porphyra.rutgers.edu/Rossoni_et_al_2019.zip | – | – |
| Galdieria sulphuraria Azora | Institute of Plant Biochemistry, HHU,Germany | http://porphyra.rutgers.edu/Rossoni_et_al_2019.zip | – | – |
| Galdieria sulphurariaMS1 | Institute of Plant Biochemistry, HHU,Germany | http://porphyra.rutgers.edu/Rossoni_et_al_2019.zip | – | – |
| Galdieria sulphurariaMtSh | Institute of Plant Biochemistry, HHU,Germany | http://porphyra.rutgers.edu/Rossoni_et_al_2019.zip | – | – |
| Galdieria sulphuraria RT22 | Institute of Plant Biochemistry, HHU,Germany | http://porphyra.rutgers.edu/Rossoni_et_al_2019.zip | – | – |
| Galdieria sulphuraria SAG21 | Institute of Plant Biochemistry, HHU,Germany | http://porphyra.rutgers.edu/Rossoni_et_al_2019.zip | – | – |
| Galdieria sulphuraria YNP5587_1 | Institute of Plant Biochemistry, HHU,Germany | http://porphyra.rutgers.edu/Rossoni_et_al_2019.zip | – | – |
| Cyanidioschyzonmerolae Soos | Institute of Plant Biochemistry, HHU,Germany | http://porphyra.rutgers.edu/Rossoni_et_al_2019.zip | – | – |
| Cyanidioschyzon merolae 10D | Refseq | ftp://ftp.ncbi.nlm.nih.gov/genomes/refseq/plant/Cyanidioschyzon_merolae/latest_assembly_versions/GCF_000091205.1_ASM9120v1 | 20.03.19 | complete genome |
| Chondrus crispus | Refseq | ftp://ftp.ncbi.nlm.nih.gov/genomes/refseq/plant/Chondrus_crispus/latest_assembly_versions/GCF_000350225.1_ASM35022v2 | 20.03.19 | scaffold |

**Suppl. Table S2:** List of all organisms that have been used in the presence-absence analysis (Figure 5). Organisms are provided with name, genome assembly accession and taxonomy. The order of organisms along the Y-axis in the presence-absence plot is according to the order used in table S2.

Provided as Excel-Table S2

**Suppl. Table S3:** Annotation of all differentially expressed transcripts of *C. reinhardtii*, *C. paradoxa* and *P. purpureum* between daylight and high-light conditions only. For a more complete annotation, only unpredicted sequences were chosen from all BLAST hits as long as they met the local identity and e-value requirements. Only transcripts with logarithmic fold-changes of at least 2 were considered (p=0.001)

Provided as Excel-Table S3

**Suppl. Table S4:** Full results of the Trinity and edgeR analysis for identification of differentially expressed genes between all tested conditions providing logarithmic fold changes of transcripts as well as the calculated means from biological triplicates. For a more complete annotation, only unpredicted sequences were chosen from all BLAST hits as long as they met the local identity and e-value requirements. Only transcripts with logarithmic fold-changes of at least 2 were considered (p=0.001 if not stated otherwise)

Provided as Excel-Table S4

**Suppl. Table S5:** List of all 112 Refseq plant and algal genomes that were used to annotate the transcripts of *C. reinhardtii*, *C. paradoxa* and *P. purpureum*.

| Organism name | Assembly accession | TaxonomyID | Assembly level | Genome release date | Downloaded |
|---|---|---|---|---|---|
| Citrus sinensis | GCF_000317415.1 | 2711 | Chromosome | 12.12.12 | 28.06.19 |
| Physcomitrella patens | GCF_000002425.4 | 3218 | Chromosome | 24.01.18 | 28.06.19 |
| Papaver somniferum | GCF_003573695.1 | 3469 | Chromosome | 18.09.18 | 28.06.19 |
| Gossypium hirsutum | GCF_000987745.1 | 3635 | Chromosome | 05.05.15 | 28.06.19 |
| Theobroma cacao | GCF_000208745.1 | 3641 | Chromosome | 09.07.16 | 28.06.19 |
| Cucumis sativus | GCF_000004075.2 | 3659 | Chromosome | 27.10.14 | 28.06.19 |
| Cucurbita pepo subsp. pepo | GCF_002806865.1 | 3664 | Chromosome | 07.12.17 | 28.06.19 |
| Populus trichocarpa | GCF_000002775.4 | 3694 | Chromosome | 24.01.18 | 28.06.19 |
| Arabidopsis thaliana | GCF_000001735.4 | 3702 | Chromosome | 15.03.18 | 28.06.19 |
| Brassica napus | GCF_000686985.2 | 3708 | Chromosome | 15.09.17 | 28.06.19 |
| Brassica rapa | GCF_000309985.1 | 3711 | Chromosome | 07.11.12 | 28.06.19 |
| Brassica oleracea var. oleracea | GCF_000695525.1 | 109376 | Chromosome | 27.05.14 | 28.06.19 |
| Malus domestica | GCF_002114115.1 | 3750 | Chromosome | 03.05.17 | 28.06.19 |
| Prunus persica | GCF_000346465.2 | 3760 | Chromosome | 02.02.17 | 28.06.19 |
| Arachis hypogaea | GCF_003086295.2 | 3818 | Chromosome | 30.05.18 | 28.06.19 |
| Cajanus cajan | GCF_000340665.1 | 3821 | Chromosome | 29.09.16 | 28.06.19 |
| Cicer arietinum | GCF_000331145.1 | 3827 | Chromosome | 16.01.13 | 28.06.19 |
| Glycine max | GCF_000004515.5 | 3847 | Chromosome | 24.07.18 | 28.06.19 |
| Glycine soja | GCF_004193775.1 | 3848 | Chromosome | 21.02.19 | 28.06.19 |
| Lupinus angustifolius | GCF_001865875.1 | 3871 | Chromosome | 16.11.16 | 28.06.19 |
| Medicago truncatula | GCF_000219495.3 | 3880 | Chromosome | 18.06.14 | 28.06.19 |
| Phaseolus vulgaris | GCF_000499845.1 | 3885 | Chromosome | 29.11.13 | 28.06.19 |
| Vigna angularis | GCF_001190045.1 | 3914 | Chromosome | 31.07.15 | 28.06.19 |
| Vigna unguiculata | GCF_004118075.1 | 3917 | Chromosome | 30.01.19 | 28.06.19 |
| Manihot esculenta | GCF_001659605.1 | 3983 | Chromosome | 10.06.16 | 28.06.19 |
| Daucus carota subsp. sativus | GCF_001625215.1 | 79200 | Chromosome | 06.05.16 | 28.06.19 |
| Capsicum annuum | GCF_000710875.1 | 4072 | Chromosome | 11.03.15 | 28.06.19 |
| Solanum lycopersicum | GCF_000188115.4 | 4081 | Chromosome | 18.04.18 | 28.06.19 |
| Olea europaea var. sylvestris | GCF_002742605.1 | 158386 | Chromosome | 03.11.17 | 28.06.19 |
| Sesamum indicum | GCF_000512975.1 | 4182 | Chromosome | 06.01.14 | 28.06.19 |
| Helianthus annuus | GCF_002127325.1 | 4232 | Chromosome | 12.05.17 | 28.06.19 |
| Cynara cardunculus var. scolymus | GCF_001531365.1 | 59895 | Chromosome | 17.04.18 | 28.06.19 |
| Oryza sativa Japonica Group | GCF_001433935.1 | 39947 | Chromosome | 10.10.15 | 28.06.19 |
| Oryza brachyantha | GCF_000231095.1 | 4533 | Chromosome | 19.01.12 | 28.06.19 |
| Setaria italica | GCF_000263155.2 | 4555 | Chromosome | 30.10.15 | 28.06.19 |
| Sorghum bicolor | GCF_000003195.3 | 4558 | Chromosome | 07.04.17 | 28.06.19 |
| Zea mays | GCF_000005005.2 | 4577 | Chromosome | 07.02.17 | 28.06.19 |
| Ananas comosus | GCF_001540865.1 | 4615 | Chromosome | 14.03.16 | 28.06.19 |
| Musa acuminata subsp. malaccensis | GCF_000313855.2 | 214687 | Chromosome | 14.11.12 | 28.06.19 |
| Asparagus officinalis | GCF_001876935.1 | 4686 | Chromosome | 06.02.17 | 28.06.19 |
| Coffea arabica | GCF_003713225.1 | 13443 | Chromosome | 08.11.18 | 28.06.19 |
| Brachypodium distachyon | GCF_000005505.3 | 15368 | Chromosome | 24.01.18 | 28.06.19 |
| Solanum pennellii | GCF_001406875.1 | 28526 | Chromosome | 11.07.14 | 28.06.19 |
| Gossypium arboreum | GCF_000612285.1 | 29729 | Chromosome | 28.09.15 | 28.06.19 |
| Gossypium raimondii | GCF_000327365.1 | 29730 | Chromosome | 20.12.12 | 28.06.19 |
| Vitis vinifera | GCF_000003745.3 | 29760 | Chromosome | 07.12.09 | 28.06.19 |
| Coffea eugenioides | GCF_003713205.1 | 49369 | Chromosome | 08.11.18 | 28.06.19 |
| Nicotiana attenuata | GCF_001879085.1 | 49451 | Chromosome | 15.11.16 | 28.06.19 |
| Elaeis guineensis | GCF_000442705.1 | 51953 | Chromosome | 13.08.13 | 28.06.19 |
| Fragaria vesca subsp. vesca | GCF_000184155.1 | 101020 | Chromosome | 24.02.11 | 28.06.19 |
| Ostreococcus tauri | GCF_000214015.3 | 70448 | Chromosome | 02.10.14 | 28.06.19 |
| Rosa chinensis | GCF_002994745.1 | 74649 | Chromosome | 15.01.19 | 28.06.19 |
| Camelina sativa | GCF_000633955.1 | 90675 | Chromosome | 24.04.14 | 28.06.19 |
| Prunus mume | GCF_000346735.1 | 102107 | Chromosome | 28.02.14 | 28.06.19 |
| Arachis duranensis | GCF_000817695.2 | 130453 | Chromosome | 25.04.17 | 28.06.19 |
| Arachis ipaensis | GCF_000816755.2 | 130454 | Chromosome | 25.04.17 | 28.06.19 |
| Vigna radiata var. radiata | GCF_000741045.1 | 3916 | Chromosome | 28.10.15 | 28.06.19 |
| Beta vulgaris subsp. vulgaris | GCF_000511025.2 | 3555 | Chromosome | 07.07.15 | 28.06.19 |
| Panicum hallii | GCF_002211085.1 | 206008 | Chromosome | 01.05.18 | 28.06.19 |
| Ziziphus jujuba | GCF_000826755.1 | 326968 | Chromosome | 06.02.15 | 28.06.19 |
| Cyanidioschyzon merolae strain 10D | GCF_000091205.1 | 280699 | Complete Genor | 11.07.07 | 28.06.19 |
| Micromonas commoda | GCF_000090985.2 | 296587 | Complete Genor | 10.04.09 | 28.06.19 |
| Aegilops tauschii subsp. tauschii | GCF_001957025.1 | 169297 | Contig | 19.01.17 | 28.06.19 |
| Prosopis alba | GCF_004799145.1 | 207710 | Contig | 17.04.19 | 28.06.19 |
| Coccomyxa subellipsoidea C-169 | GCF_000258705.1 | 574566 | Contig | 13.04.12 | 28.06.19 |
| Chondrus crispus | GCF_000350225.1 | 2769 | Scaffold | 22.05.13 | 28.06.19 |
| Chlamydomonas reinhardtii | GCF_000002595.1 | 3055 | Scaffold | 15.10.07 | 28.06.19 |
| Volvox carteri f. nagariensis | GCF_000143455.1 | 3068 | Scaffold | 08.07.10 | 28.06.19 |
| Spinacia oleracea | GCF_002007265.1 | 3562 | Scaffold | 27.02.17 | 28.06.19 |
| Carica papaya | GCF_000150535.2 | 3649 | Scaffold | 06.05.08 | 28.06.19 |
| Cucumis melo | GCF_000313045.1 | 3656 | Scaffold | 05.10.12 | 28.06.19 |
| Cucurbita maxima | GCF_002738345.1 | 3661 | Scaffold | 31.10.17 | 28.06.19 |
| Cucurbita moschata | GCF_002738365.1 | 3662 | Scaffold | 31.10.17 | 28.06.19 |
| Momordica charantia | GCF_001995035.1 | 3673 | Scaffold | 26.12.16 | 28.06.19 |
| Raphanus sativus | GCF_000801105.1 | 3726 | Scaffold | 29.09.15 | 28.06.19 |
| Abrus precatorius | GCF_003935025.1 | 3816 | Scaffold | 11.12.18 | 28.06.19 |
| Hevea brasiliensis | GCF_001654055.1 | 3981 | Scaffold | 01.06.16 | 28.06.19 |
| Ricinus communis | GCF_000151685.1 | 3988 | Scaffold | 07.07.11 | 28.06.19 |
| Nicotiana sylvestris | GCF_000393655.1 | 4096 | Scaffold | 16.05.13 | 28.06.19 |
| Nicotiana tabacum | GCF_000715135.1 | 4097 | Scaffold | 29.05.14 | 28.06.19 |
| Nicotiana tomentosiformis | GCF_000390325.2 | 4098 | Scaffold | 16.05.13 | 28.06.19 |
| Solanum tuberosum | GCF_000226075.1 | 4113 | Scaffold | 19.09.11 | 28.06.19 |
| Erythranthe guttata | GCF_000504015.1 | 4155 | Scaffold | 02.04.14 | 28.06.19 |
| Lactuca sativa | GCF_002870075.1 | 4236 | Scaffold | 09.01.18 | 28.06.19 |
| Nelumbo nucifera | GCF_000365185.1 | 4432 | Scaffold | 08.08.13 | 28.06.19 |
| Camellia sinensis | GCF_004153795.1 | 4442 | Scaffold | 11.02.19 | 28.06.19 |
| Amborella trichopoda | GCF_000471905.2 | 13333 | Scaffold | 30.09.13 | 28.06.19 |
| Tarenaya hassleriana | GCF_000463585.1 | 28532 | Scaffold | 05.09.13 | 28.06.19 |
| Ipomoea nil | GCF_001879475.1 | 35883 | Scaffold | 01.09.16 | 28.06.19 |
| Micromonas pusilla CCMP1545 | GCF_000151265.2 | 564608 | Scaffold | 09.04.09 | 28.06.19 |
| Prunus avium | GCF_002207925.1 | 42229 | Scaffold | 12.06.17 | 28.06.19 |
| Phoenix dactylifera | GCF_000413155.1 | 42345 | Scaffold | 24.06.13 | 28.06.19 |
| Juglans regia | GCF_001411555.1 | 51240 | Scaffold | 22.10.15 | 28.06.19 |
| Quercus suber | GCF_002906115.1 | 58331 | Scaffold | 29.01.18 | 28.06.19 |
| Arabidopsis lyrata subsp. lyrata | GCF_000004255.2 | 81972 | Scaffold | 26.11.16 | 28.06.19 |
| Chenopodium quinoa | GCF_001683475.1 | 63459 | Scaffold | 19.05.17 | 28.06.19 |
| Durio zibethinus | GCF_002303985.1 | 66656 | Scaffold | 15.09.17 | 28.06.19 |
| Eucalyptus grandis | GCF_000612305.1 | 71139 | Scaffold | 02.05.14 | 28.06.19 |
| Eutrema salsugineum | GCF_000478725.1 | 72664 | Scaffold | 05.11.13 | 28.06.19 |
| Populus euphratica | GCF_000495115.1 | 75702 | Scaffold | 12.08.14 | 28.06.19 |
| Phalaenopsis equestris | GCF_001263595.1 | 78828 | Scaffold | 07.08.15 | 28.06.19 |
| Capsella rubella | GCF_000375325.1 | 81985 | Scaffold | 23.04.13 | 28.06.19 |
| Citrus clementina | GCF_000493195.1 | 85681 | Scaffold | 08.11.13 | 28.06.19 |
| Selaginella moellendorffii | GCF_000143415.4 | 88036 | Scaffold | 06.07.10 | 28.06.19 |
| Herrania umbratica | GCF_002168275.1 | 108875 | Scaffold | 05.06.17 | 28.06.19 |
| Galdieria sulphuraria | GCF_000341285.1 | 130081 | Scaffold | 25.02.13 | 28.06.19 |
| Monoraphidium neglectum | GCF_000611645.1 | 145388 | Scaffold | 26.02.15 | 28.06.19 |
| Jatropha curcas | GCF_000696525.1 | 180498 | Scaffold | 02.06.14 | 28.06.19 |
| Pyrus xbretschneideri | GCF_000315295.1 | 225117 | Scaffold | 03.12.12 | 28.06.19 |
| Chlorella variabilis | GCF_000147415.1 | 554065 | Scaffold | 16.09.10 | 28.06.19 |
| Dendrobium catenatum | GCF_001605985.2 | 906689 | Scaffold | 12.12.17 | 28.06.19 |
| Morus notabilis | GCF_000414095.1 | 981085 | Scaffold | 07.08.13 | 28.06.19 |
